# Inhibitory subpopulations in preBötzinger Complex play distinct roles in modulating inspiratory rhythm and pattern

**DOI:** 10.1101/2023.08.07.552303

**Authors:** Zheng Chang, Jordan Skach, Kaiwen Kam

## Abstract

Inhibitory neurons embedded within mammalian neural circuits shape breathing, walking, chewing, and other rhythmic motor behaviors. At the core of the neural circuit controlling breathing is the preBötzinger Complex (preBötC), a nucleus in the ventrolateral medulla necessary for generation of inspiratory rhythm. In the preBötC, a recurrently connected network of glutamatergic Dbx1-derived (Dbx1^+^) neurons generates rhythmic inspiratory drive. Functionally and anatomically intercalated among preBötC Dbx1^+^ neurons are GABAergic (GAD1/2^+^) and glycinergic (GlyT2^+^) neurons, whose roles in breathing remain unclear. To elucidate the inhibitory microcircuits within preBötC, we first characterized the spatial distribution of molecularly-defined preBötC inhibitory subpopulations in double reporter mice expressing either the red fluorescent protein tdTomato or EGFP in GlyT2^+^, GAD1^+^, or GAD2^+^ neurons. We found that, in neonatal mice, the majority of preBötC inhibitory neurons expressed a combination of GlyT2 and GAD2 while a much smaller subpopulation also expressed GAD1. To determine the functional role of these subpopulations, we used holographic photostimulation, a patterned illumination technique with high spatiotemporal resolution, in rhythmically active medullary slices from neonatal Dbx1^tdTomato^;GlyT2^EGFP^ and Dbx1^tdTomato^;GAD1^EGFP^ double reporter mice. Stimulation of 4 or 8 preBötC GlyT2^+^ neurons during endogenous rhythm prolonged the interburst interval in a phase-dependent manner and increased the latency to burst initiation when bursts were evoked by stimulation of Dbx1^+^ neurons. In contrast, stimulation of 4 or 8 preBötC GAD1^+^ neurons did not affect interburst interval or latency to burst initiation. Instead, photoactivation of GAD1^+^ neurons during the inspiratory burst prolonged endogenous and evoked burst duration and decreased evoked burst amplitude. We conclude that the majority of preBötC inhibitory neurons express both GlyT2 and GAD2 and modulate breathing rhythm by delaying burst initiation while a smaller GAD1^+^ subpopulation shapes inspiratory patterning by altering burst duration and amplitude.

## Introduction

Breathing, walking, and other rhythmic motor behaviors are controlled by networks of excitatory and inhibitory neurons that coordinate and adjust muscle activity to meet physiological demand. At the core of the neural circuit controlling breathing is the preBötzinger Complex (preBötC), a compact nucleus in the ventrolateral medulla that generates breathing rhythm and is hypothesized to organize other rhythmic orofacial movements (Ashhad et al., 2022; Moore et al., 2014; Smith et al., 1991). In the preBötC, a recurrently connected network of glutamatergic Dbx1-derived (Dbx1^+^) neurons generates rhythmic inspiratory drive to inspiratory premotoneurons and motoneurons (Ashhad et al., 2022; Bouvier et al., 2010; Gray et al., 2010; Koizumi et al., 2008). Also within the preBötC are neurons that are glycinergic (defined by expression of the vesicular glycine transporter GlyT2; GlyT2^+^; Koizumi et al., 2013; Morgado-Valle et al., 2010; Winter et al., 2009) and GABAergic (defined by expression of either or both of the two isoforms of glutamic acid decarboxylase, GAD1/GAD67 and GAD2/GAD65; GAD1/2^+^; Erlander et al., 1991; Koizumi et al., 2013; Kuwana et al., 2006). GlyT2^+^ and GAD1/2^+^ neurons are functionally and anatomically intercalated among Dbx1^+^ neurons, with many, but not all, showing inspiratory-modulated activity (Koizumi et al., 2013; Kuwana et al., 2006; Morgado-Valle et al., 2010; Winter et al., 2009). Co-expression of GlyT2 with either GAD2 or GAD1 in preBötC and thus co-release of glycine and GABA is observed postnatally (Hirrlinger et al., 2019; Koizumi et al., 2013; Oke et al., 2023; Rahman et al., 2013). However, a developmental switch leads to many neurons only expressing either GlyT2 or GAD2 in adult animals (Hirrlinger et al., 2019; Oke et al., 2023).

Inhibition regulates many aspects of breathing (Baertsch et al., 2018; Dutschmann and Dick, 2012; Ezure and Tanaka, 2004; Ezure et al., 2003; Janczewski et al., 2013; Kubin et al., 2006; Shao and Feldman, 1997; Sherman et al., 2015; Smith et al., 2007) and plays a role in the etiology of neurodevelopmental disorders with respiratory phenotypes, such as Rett Syndrome (Abdala et al., 2010; Chao et al., 2010). Pharmacological disinhibition of the preBötC and a nearby respiratory-related nucleus, the Bötzinger Complex (BötC), in *in situ* preparations that generate a motor rhythm with distinct inspiratory, postinspiratory, and expiratory phases causes inspiratory and postinspiratory phases to fuse and can produce tonic activity on respiratory motor nerves (Dutschmann and Paton, 2002; Marchenko et al., 2016). Optogenetic stimulation of medullary GlyT2^+^ neurons *in vivo* delays inspiration, modulates tidal volume, and causes apneas (Ausborn et al., 2018; Fortuna et al., 2019; Hulsmann et al., 2021; Sherman et al., 2015), but has also been shown to increase inspiratory frequency by reducing excitability during inspiration, shortening postinspiratory refractoriness, and evoking post-inhibitory rebound (Baertsch et al., 2018). Optogenetic photoinhibition of GlyT2^+^ neurons increases tidal volume and shortens expiratory duration (Sherman et al., 2015; Vafadari et al., 2023). preBötC GAD1^+^ neurons project ipsilaterally to downstream XII premotoneurons and motoneurons at the same coronal level in the medullary slice (Koizumi et al., 2013), indicative of a role in modulating inspiratory burst pattern as activity is transmitted from the preBötC to XII nerve. Proposed roles for preBötC inhibitory neurons thus include ensuring the alternating silence of antagonist inspiratory and expiratory muscles (Ezure et al., 2003; Janczewski et al., 2013), modulating motor nerve burst pattern (Janczewski et al., 2013; Shao and Feldman, 1997), inducing apneas for regulatory and protective reflexes and volitional behaviors (Ezure and Tanaka, 2004; Janczewski et al., 2013; Kubin et al., 2006), contributing to preBötC excitation-inhibition balance (Ashhad and Feldman, 2020; Ramirez and Baertsch, 2018), and generating rhythm (Smith et al., 2007).

There are, however, conflicting data and interpretations over the role of preBötC inhibitory neurons in breathing (Janczewski et al., 2013; Marchenko et al., 2016; Smith et al., 2007). In contrast to *in situ* data (Dutschmann and Paton, 2002; Marchenko et al., 2016), pharmacological block of inhibitory synaptic transmission *in vitro* or *in vivo* has minimal effects on respiratory rhythm generation (Janczewski et al., 2013; Shao and Feldman, 1997). Genetic deletion of GlyT2 affects, but does not eliminate, breathing in mice and does not significantly affect preBötC rhythmic activity postnatally (Hulsmann et al., 2019). Further, previous pharmacological and optogenetic manipulations targeting preBötC inhibitory neurons may also have affected unspecified numbers and types of inhibitory neurons in adjacent BötC and potentially throughout the medulla. The specific effects of activation or inhibition of preBötC GAD1/2^+^ neurons have also not been explored, and whether distinct inhibitory subpopulations mediate the myriad functions attributed to inhibition in preBötC is not known. Thus, a clear picture for the role of preBötC inhibitory neurons in breathing has remained elusive (Abdala et al., 2015; Ashhad et al., 2022; Dick et al., 2018; Feldman and Kam, 2015; Ramirez and Baertsch, 2018).

To shed light on the molecular heterogeneity and function of inhibitory neurons and microcircuits within the preBötC, we used double reporter mice where red and green fluorescent proteins were expressed in Dbx1^+^, GlyT2^+^, GAD1^+^, or GAD2^+^ neurons. We found that the majority of preBötC inhibitory neurons in the postnatal mouse express both GlyT2 and GAD2 while a much smaller subpopulation also expresses GAD1. To determine whether molecularly-defined preBötC inhibitory neurons showed any functional differences, we used holographic photostimulation, which can dynamically and selectively excite small sets of neurons within a population with exceptional spatiotemporal resolution, in rhythmic medullary slices from double reporter mice (Kam et al., 2013b; Lutz et al., 2008; Zahid et al., 2010). Holographic photostimulation of GlyT2^+^ neurons in the preinspiratory period delayed burst initiation, consistent with a role for preBötC GlyT2^+^ neurons in regulating inspiratory rhythm, whereas photostimulation of GAD1^+^ neurons during inspiration increased burst duration and decreased burst amplitude, pointing to a role for preBötC GAD1^+^ neurons in shaping inspiratory pattern. We therefore reveal functional differences among embedded preBötC inhibitory microcircuits in the clinically important motor program for breathing.

## Results

### GAD1 defines a distinct preBötC inhibitory subpopulation

We first examined the spatial distribution of neurons expressing the transcription factor Dbx1, which defines an excitatory preBötC subpopulation necessary for breathing (Bouvier et al., 2010; Gray et al., 2010), and/or the glycinergic inhibitory marker GlyT2 (Zeilhofer et al., 2005). We used P0-P4 double reporter mice of either sex expressing the red fluorescent protein tdTomato in Dbx1^+^ neurons and enhanced green fluorescent protein (EGFP) in GlyT2^+^ neurons. Double reporter mice were generated using a cre-lox strategy crossing Dbx1^cre^ (Bielle et al., 2005) with Rosa26^LoxP-Stop-LoxP-tdTomato^ (Madisen et al., 2010) in combination with GlyT2^EGFP^ BAC transgenic reporter mice (Zeilhofer et al., 2005). We counted neurons expressing tdTomato, EGFP, or both fluorescent proteins within a 150 μm radius hemi-cylinder ventral to nucleus ambiguus that extended rostrocaudally from 360 to 480 μm caudal to the caudal pole of facial motor nucleus, a region corresponding anatomically to preBötC (Ruangkittisakul et al., 2014). Within this region, there were similar numbers of Dbx1^+^ and GlyT2^+^ neurons (Dbx1^+^: 662.8 ± 189.9; GlyT2^+^: 636.3 ± 132.8; n=4; Figure 1A-C). Co-expression of Dbx1 and GlyT2 in this region was observed (175.3 ± 62.5; n=4; Figure 1A-C), constituting 26.4% of Dbx1^+^ neurons and 27.5% of GlyT2^+^ neurons. No tdTomato labeling was detected in preBötC from cre-negative Dbx1^+/+^;Rosa26^LoxP-Stop-LoxP-tdTomato^ animals (data not shown).

**Figure 1.**
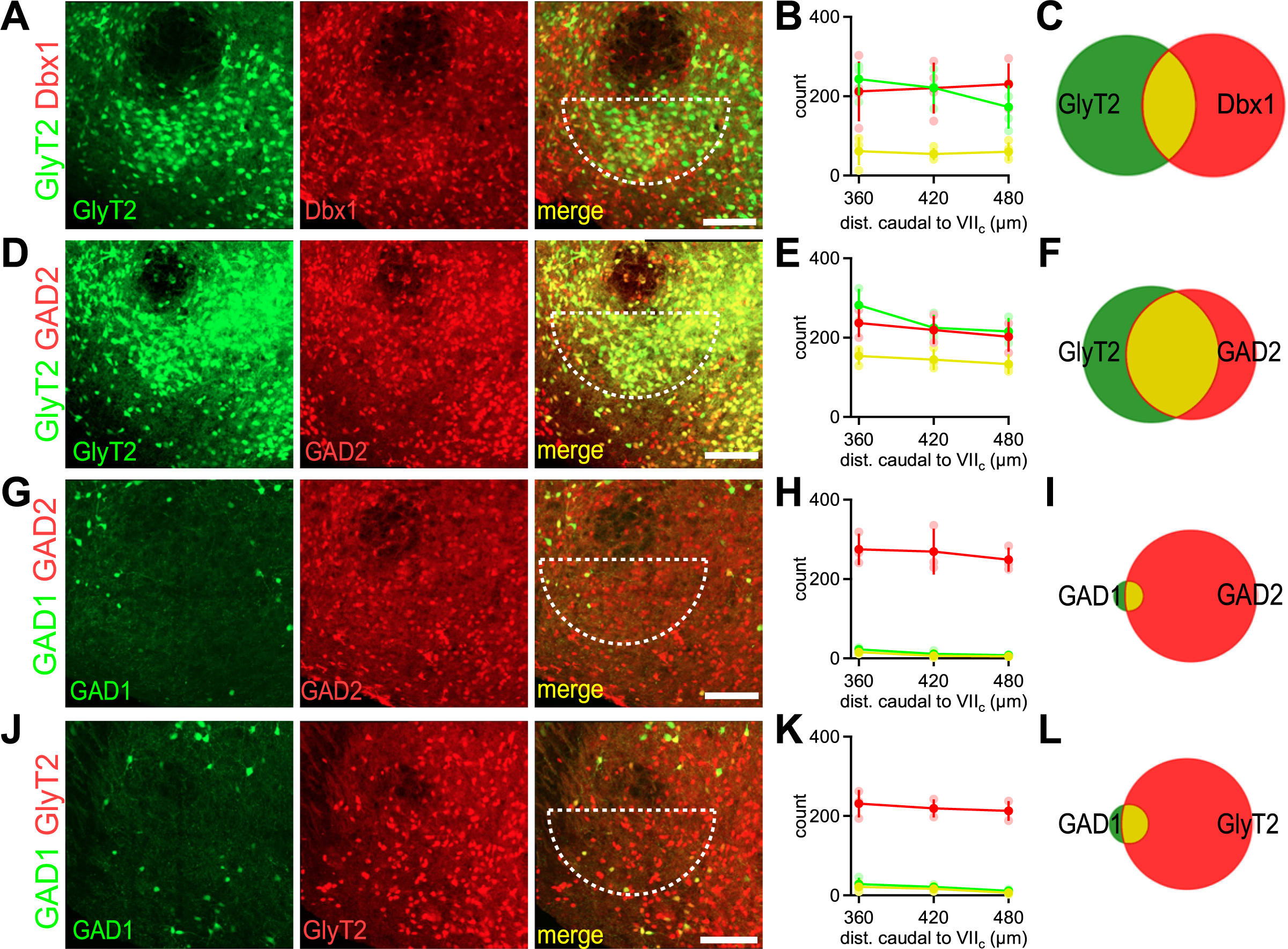
Spatial distribution of preBötC inhibitory subpopulations. Characterization of the number and spatial distribution of molecularly-defined preBötC neurons from **(A-C)** Dbx1^tdTomato^;GlyT2^EGFP^, **(D-F)** GAD2^tdTomato^;GlyT2^EGFP^, **(G-I)** GAD2^tdTomato^;GAD1^EGFP^, and **(J-L)** GlyT2^tdTomato^;GAD1^EGFP^ mice. **(A, D, G, J)** Confocal z-projection of EGFP, tdTomato, and merged channels from a 60 µm section from the preBötC of the specified genotype. The dotted semicircle represents the area where neurons are counted. Scale bar, 100 µm. **(B, E, H, K)** Rostrocaudal distributions of EGFP (green), tdTomato (red), and co-expressing (yellow) neurons. Counts are the sum of neurons from left and right preBötC from 60 µm sections at the specified level, defined relative to the caudal pole of facial motor nucleus (VII_c_). **(C, F, I, L)** Venn diagrams representing the average total counts of preBötC EGFP (green), tdTomato (red), and co-expressing (yellow) neurons. Total counts are the sum of neurons from left and right preBötC across three 60 µm sections that span the 360-480 µm rostrocaudal extent of preBötC.

To determine the molecular phenotype of preBötC inhibitory neurons, we characterized the pairwise spatial distribution and levels of co-expression of inhibitory markers in the preBötC of P0-P4 transgenic double reporter mice expressing either tdTomato or EGFP in GlyT2^+^, GAD1^+^, or GAD2^+^ neurons. Double reporter mice were generated using a cre-lox strategy, crossing GlyT2^cre^ (Foster et al., 2015) or GAD2^cre^ (Taniguchi et al., 2011) with Rosa26^LoxP-Stop-LoxP-tdTomato^ in combination with GlyT2^EGFP^ or GAD1^EGFP^ BAC transgenic reporter mice (Chattopadhyaya et al., 2004; Zeilhofer et al., 2005).

In GAD2tdTomato_;GlyT2_EGFP _mice (GAD2_cre _x Rosa26_LoxP-Stop-LoxP-tdTomato _x GlyT2_EGFP_), GlyT2_+ and GAD2^+^ neurons were found in approximately equal numbers with GlyT2^+^ slightly outnumbering GAD2^+^ (GlyT2^+^: 722 ± 106.7; GAD2^+^: 658.3 ± 107.1; n=3; Figure 1D-F). Co-expression of GlyT2 and GAD2 was high (GlyT2^+^/GAD2^+^: 431 ± 66; n=3) with 59.7% of GlyT2^+^ expressing GAD2 and 65.5% of GAD2^+^ expressing GlyT2 (Figure 1D-F). In contrast to the high density of GAD2^+^ and GlyT2^+^ preBötC neurons, significantly fewer GAD1^+^ neurons were observed in preBötC. In GAD2^tdTomato^;GAD1^EGFP^ mice (GAD2^cre^ x Rosa26^LoxP-Stop-LoxP-tdTomato^ x GAD1^EGFP^) mice, the number of GAD1^+^ neurons was much lower than GAD2^+^ neurons in preBötC (GAD2^+^: 792 ± 125.4; GAD1^+^: 41 ± 13.9; n=3; Figure 1G-I). Most GAD1^+^ neurons expressed GAD2 (GAD2^+^/GAD1^+^: 26.3 ± 10.4; n=3) with 64.3% of GAD1^+^ neurons expressing GAD2, but only 3.3% of GAD2^+^ neurons expressing GAD1 (Figure 1G-I). A similar distribution was observed in GlyT2^tdTomato^;GAD1^EGFP^ mice (GlyT2^cre^ x Rosa26^LoxP-Stop-LoxP-tdTomato^ x GAD1^EGFP^). In these mice, the number of GAD1^+^ neurons was much lower than GlyT2^+^ neurons (GlyT2^+^: 662 ± 81.1; GAD1^+^: 60.3 ± 24; n=3; Figure 1J-L). Most GAD1^+^ neurons expressed GlyT2 (GlyT2^+^/GAD1^+^: 43.3 ± 18.5; n=3) with 71.8% of GAD1^+^ neurons expressing GlyT2, but only 6.5% of GlyT2^+^ neurons expressing GAD1 (Figure 1J-L). Across all genotypes, the number of preBötC GAD1^+^ neurons (50.7 ± 20.5; n=6) was significantly lower than the number of preBötC GlyT2^+^ (669.7 ± 106.3; n=10), GAD2^+^ (729.7 ± 136.5; n=6), and Dbx1^+^ (662.8 ± 189.9; n=4) neurons (One-way ANOVA, F(3,22)=45.2, p=1.4 × 10^-9^; post-hoc Tukey test, GAD1^+^ vs GlyT2^+^: p=5.1 × 10^-9^; GAD1^+^ vs GAD2^+^: p=7.2 × 10^-9^; GAD1^+^ vs Dbx1^+^: p=3.0 × 10^-7^). Thus, we found that inhibitory neurons were heterogeneous in their expression of these molecular markers and suggest that two major subpopulations of inhibitory neurons in the preBötC of neonatal mice are GAD2^+^/GlyT2^+^ co-expressing neurons and GAD1^+^ neurons that likely also express GAD2 and/or GlyT2.

### Threshold preBötC GlyT2^+^ stimulation delays burst initiation

We sought to examine whether these inhibitory preBötC subpopulations had distinct functional roles in inspiratory rhythm and pattern generation. A major issue with functional studies involving preBötC inhibitory neurons *in vitro* and *in vivo* is the difficulty in restricting perturbations to preBötC neurons due to the high density of inhibitory neurons in nearby BötC and numerous inhibitory afferents terminating in preBötC from other medullary sites. To selectively target preBötC GlyT2^+^ neurons and assess their functional role, we used holographic photostimulation in rhythmically active medullary slices obtained from neonatal Dbx1^tdTomato^;GlyT2^EGFP^ (Dbx1^cre^ x Rosa26^LoxP-Stop-LoxP-tdTomato^ x GlyT2^EGFP^) double reporter mice (Kam et al., 2013b; Lutz et al., 2008; Zahid et al., 2010). Glutamate uncaging of a 10 μm soma-centered spot over a single preBötC GlyT2^+^ neuron adjacent to Dbx1^+^ neurons resulted in a burst of action potentials (Figure 2A), similar to the responses of Dbx1^+^ neurons to photostimulation (Kam et al., 2013b; Sun et al., 2019).

**Figure 2.**
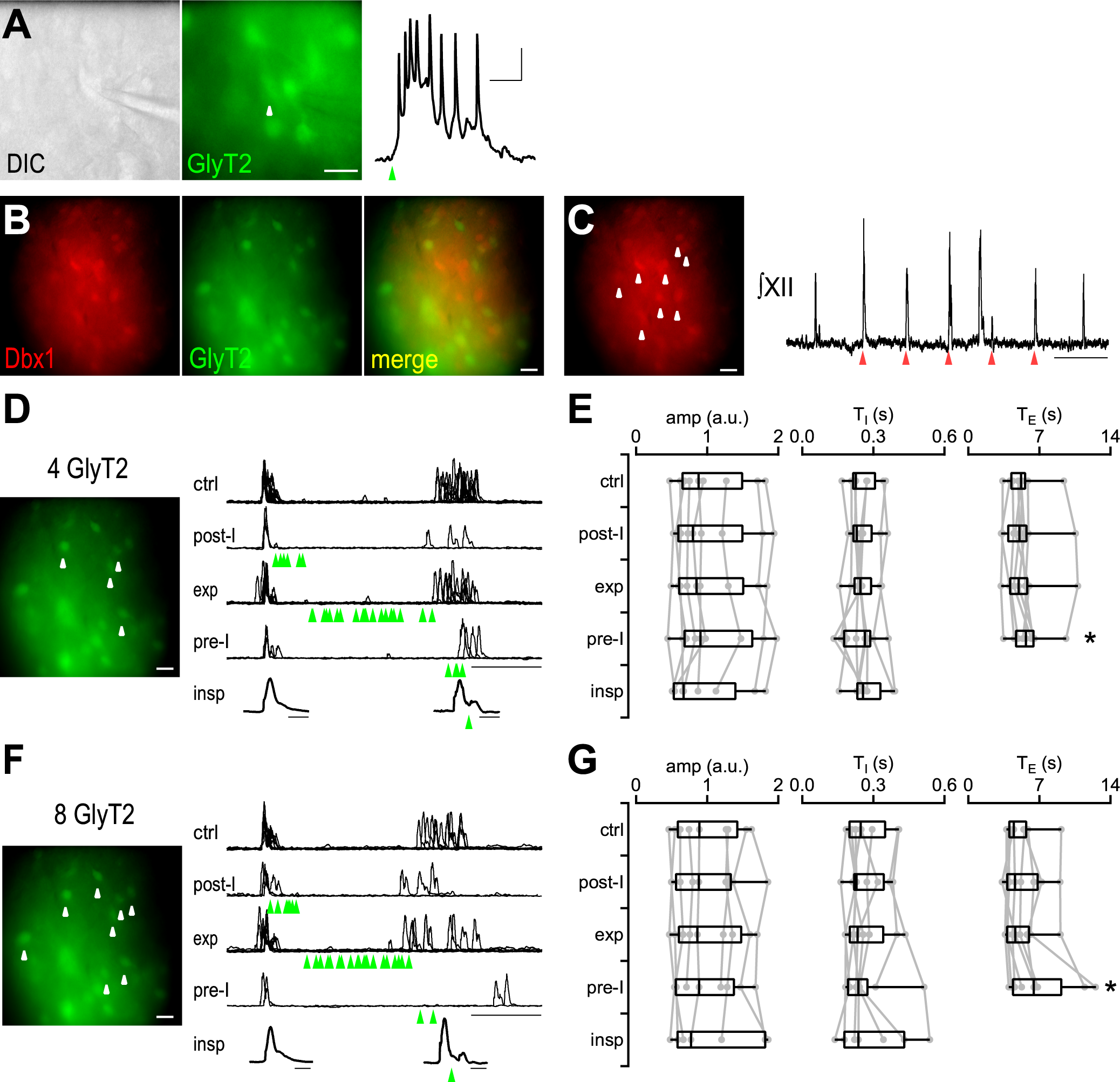
Stimulation of preBötC GlyT2^+^ neurons during endogenous rhythm increases T_E_. **(A)** DIC and fluorescence images of a Dbx1^tdTomato^;GlyT2^EGFP^ mouse. Whole cell current clamp response of a GlyT2^+^ neuron (white arrow) to photostimulation (green arrow). Scale bar, 10 mV, 100 ms. **(B)** Fluorescent images of preBötC from Dbx1^tdTomato^;GlyT2^EGFP^ mouse. Scale bar, 100 µm. **(C)** *left*, Fluorescent tdTomato image of preBötC. White arrows mark targeted neurons. Scale bar, 100 µm. *right*, Integrated XII (∫XII) trace shows response to photostimulation of 8 Dbx1^+^ neurons. Scale bar, 5 s. **(D)** *left*, Fluorescent image showing 4 preBötC GlyT2^+^ neurons targeted for photostimulation (white arrows). Scale bar, 100 µm. *right*, Overlay of cycles of ∫XII activity under control conditions with no stimulation and when stimulation occurred during post-inspiratory (post-I), expiratory (exp), and preinspiratory (pre-I) phases. Green arrows mark time of stimulation. Scale bar, 2 s. Average control inspiratory burst shape (*bottom left*) and burst shape when stimulation occurred (*bottom right*) during the inspiratory burst (insp). Green arrow marks time of stimulation. Scale bar, 0.2 s. **(E)** Grouped amplitude (amp), burst duration (T_I_), and interburst interval (T_E_) for cycles with no stimulation (ctrl), and when stimulation of 4 GlyT2^+^ preBötC neurons occurred during post-I, exp, and pre-I. T_I_ and amp are also shown when stimulation occurred during bursts (insp). *, p<0.05, Friedman test, post-hoc Wilcoxon signed rank test with Bonferroni-Holm correction, n=9. (**F)** *left*, Fluorescent image showing 8 preBötC GlyT2^+^ neurons targeted for photostimulation (white arrows). Scale bar, 100 µm. *right*, Overlay of cycles of ∫XII activity under control conditions with no stimulation, and when stimulation occurred during post-inspiratory (post-I), expiratory (exp), and preinspiratory (pre-I) phases. Green arrows mark time of stimulation. Scale bar, 2 s. Average control inspiratory burst shape (*bottom left*) and burst shape when stimulation occurred (*bottom right*) during the inspiratory burst (insp). Green arrow marks time of stimulation. Scale bar, 0.2 s. **(G)** Grouped amplitude (amp), burst duration (T_I_), and interburst interval (T_E_) for cycles with no stimulation (ctrl), and when stimulation of 8 GlyT2^+^ preBötC neurons occurred during post-I, exp, and pre-I. T_I_ and amp are also shown when stimulation occurred during bursts (insp). *, p<0.05, Friedman test, post-hoc Wilcoxon signed rank test with Bonferroni-Holm correction, n=9.

To confirm that targeted neurons were in preBötC, we examined Dbx1/GlyT2 colocalization, which is minimal in preBötC, and whether Dbx1^+^ neuron stimulation could evoke bursts and entrain rhythm, a property of preBötC Dbx1^+^ neurons (Bouvier et al., 2010; Gray et al., 2010; Kam et al., 2013b; Sun et al., 2019). Consistent with the Dbx1^+^ neurons being in preBötC, we observed no colocalization of Dbx1^+^ and GlyT2^+^ in our stimulation field (Figure 2B). We then photostimulated Dbx1^+^ neurons and calculated a probability of stimulating a burst (P_stim_) by dividing the number of successfully evoked bursts by the number of laser stimuli. Holographic photostimulation of 8 Dbx1^+^ neurons at intervals slightly shorter than the endogenous rhythm resulted in generation of inspiratory bursts in a median of 94% [IQR 76-95%] of stimuli (n=9; Figure 2C), consistent with these neurons playing a role in inspiratory rhythmogenesis.

In the same field, we selected 4 or 8 GlyT2^+^ neurons for low frequency stimulation, which would allow sampling of all phases of endogenous rhythm. The interval between stimuli was greater than 2 times the average period to allow for unstimulated cycles in between stimulated cycles. The phase of stimulation was defined as the time of stimulation after a burst divided by the average interburst interval (T_E_) of unstimulated cycles (control T_E_). We categorized the calculated phases of stimulation into those occurring during the postinspiratory period (post-I; <20% of control T_E_ immediately following a burst), during expiration (Exp; 20-90% of control T_E_ after post-I), during the preinspiratory period (pre-I; 90-110% of control T_E_ following Exp if stimulation did not occur during a burst), and during inspiration (stimulation occurring during a burst) and measured the effect on T_E_.

Photostimulation of 4 preBötC GlyT2^+^ neurons adjacent to the photostimulated Dbx1^+^ neurons during post-I and Exp did not affect T_E_, but pre-I stimulation resulted in a 9.2% increase in T_E_ (5.67 s [IQR 4.71-6.42 s]) compared to control (5.19 s [IQR 4.23-5.57 s]) that was statistically significant (Friedman test, χ^2^(3)=10.2, p=0.02; post-hoc Wilcoxon signed rank test with Bonferroni-Holm correction, ctrl vs. post-I: p=0.6; ctrl vs. Exp: p=1; ctrl vs. pre-I: p=0.004; n=9; Figure 2D, E). Amplitude (amp) and duration (T_I_) of the burst subsequent to photostimulation of 4 GlyT2^+^ neurons was not significantly different from control bursts (Friedman test, amp: χ^2^(3)=3.3, p=0.35; T_I_: χ^2^(3)=0.3, p=0.95; n=9; Figure 2D, E). While stimulation of 8 GlyT2^+^ neurons during post-I and Exp also did not affect T_E_, stimulation during pre-I produced a significant 43.8% increase in T_E_ (6.44 s [IQR 4.40-9.15 s]) compared to control (4.48 s [IQR 4.04-5.71 s]; Friedman test, χ^2^(3)=14.1, p=0.003; post-hoc Wilcoxon signed rank test with Bonferroni-Holm correction, ctrl vs. post-I: p=1; ctrl vs. Exp: p=0.7; ctrl vs. pre-I: p=0.004; n=9; Figure 2F, G). Amplitude and T_I_ of the subsequent burst following stimulation of 8 GlyT2^+^ neurons at any phase were not significantly different from control (Friedman test, amp: χ^2^(3)=1.3, p=0.7; T_I_: χ^2^(3)=0.3, p=0.6; n=9; Figure 2F, G). When 4 or 8 GlyT2^+^ neurons were activated during an inspiratory burst, we observed no significant effects on inspiratory burst amplitude or duration (Wilcoxon signed rank test, 4 GlyT2^+^ amp: p=0.1; 4 GlyT2^+^ T_I_: p=0.2; 8 GlyT2^+^ amp: p=0.2; 8 GlyT2^+^ T_I_: p=0.07; n=9; Figure 2D-G).

We hypothesized that the specific effect of pre-I stimulation of 8 GlyT2^+^ neurons on T_E_ was due to preinspiration being a particularly sensitive period of increasing synchronization during rhythmogenesis (Ashhad and Feldman, 2020; Ashhad et al., 2022; Kam et al., 2013b). A similar process may be initiated by photostimulation of 4-9 preBötC neurons (Kam et al., 2013b). The latency observed when triggering synchronous activity in a small group of Dbx1^+^ neurons may represent the same assembly mechanism occurring during pre-I (Kam et al., 2013b; Sun et al., 2019). If true, stimulation of GlyT2^+^ neurons following stimulation of 8 Dbx1^+^ neurons should produce a prolonged latency to burst generation, analogous to the increased period observed when 8 GlyT2^+^ neurons were photostimulated in pre-I during endogenous rhythm. We therefore took advantage of the ability of the holographic system to display multiple patterns to stimulate Dbx1^+^ neurons and GlyT2^+^ neurons sequentially and examine how inhibition affects this assembly process. We first established a baseline by holographically photostimulating 8 Dbx1^+^ neurons at intervals just shorter than the endogenous interval. This protocol resulted in entrainment of the rhythm (P_stim_: 94.1% [IQR 89-94%]; n=9; Figure 3A, E). The latency between laser stimulation of 8 Dbx1^+^ neurons and burst initiation was 0.15 s [IQR 0.13-0.16 s] (n=9; Figure 3A, E).

**Figure 3.**
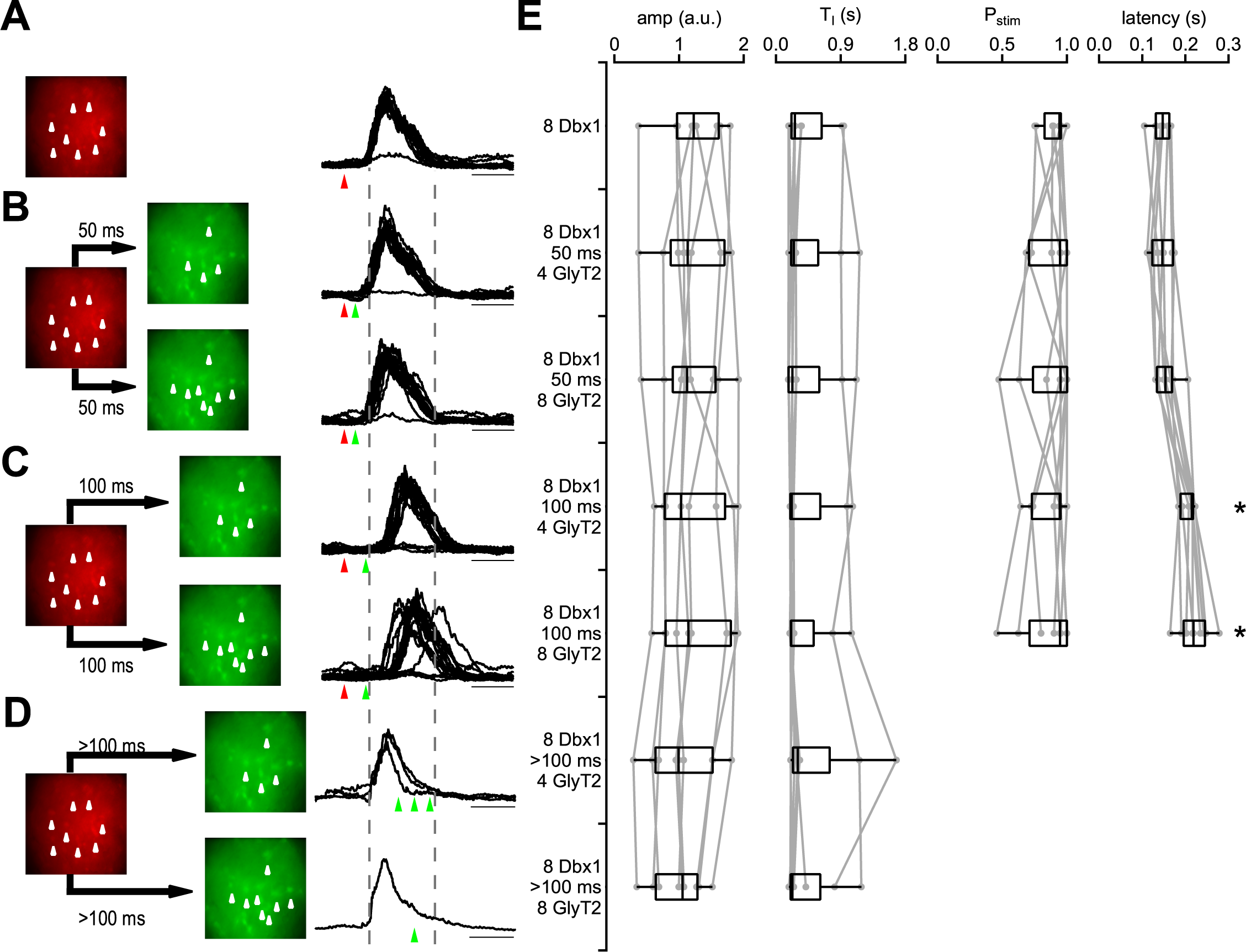
Stimulation of preBötC GlyT2^+^ neurons increases the latency to burst initiation for evoked bursts. **(A-D)** *left*, Fluorescent images of preBötC from a Dbx1^tdTomato^;GlyT2^EGFP^ mouse and, *right*, overlaid traces from the ∫XII recording of evoked bursts triggered by **(A)** 8 Dbx1^+^ neurons, **(B)** 8 Dbx1^+^ neurons followed by 4 or 8 GlyT2^+^ neurons after a 50 ms delay, **(C)** 8 Dbx1^+^ neurons followed by 4 or 8 GlyT2^+^ neurons after a 100 ms delay, and **(D)** 8 Dbx1^+^ neurons followed by 4 or 8 GlyT2^+^ neurons after a >100 ms delay (during the evoked burst). White arrows indicate targeted neurons, Red arrows indicate time of 8 Dbx1^+^ stimulation, Green arrows indicate time of 4 or 8 GlyT2^+^ stimulation 50 ms, 100, ms, and > 100 ms after Dbx1^+^ stimulation. Dotted lines indicate average beginning and end of burst evoked by 8 Dbx1^+^ neurons. Scale bar, 0.2 s. **(E)** Grouped data showing amplitude (amp), burst duration (T_I_), probability of stimulation (P_stim_), and latency to burst initiation following stimulation of Dbx1^+^ and GlyT2^+^ neurons. *, p<0.05, Friedman test, post-hoc Wilcoxon signed rank test with Bonferroni-Holm correction, n=9.

We then examined the effects of stimulating 4 or 8 GlyT2^+^ neurons 50 or 100 ms following stimulation of 8 Dbx1^+^ neurons (Figure 3B, C). While sequential stimulation of 4 or 8 GlyT2^+^ neurons 50 ms after stimulation of 8 Dbx1^+^ neurons did not significantly alter the latency preceding the burst (Figure 3B), increasing the interval between 8 Dbx1^+^ and 4 or 8 GlyT2^+^ neuron stimulation to 100 ms significantly increased the latency by 45.8% for 4 GlyT2^+^ neurons (0.22 s [IQR 0.19-0.22 s]) and by 48.6% for 8 GlyT2^+^ neurons (0.22 s [IQR 0.20-0.25 s]) compared to 8 Dbx1^+^ neuron stimulation alone (Friedman test, χ^2^(4)=26.8, p=2×10^-5^; post-hoc Wilcoxon signed rank test with Bonferroni-Holm correction, 8 Dbx1^+^/50 ms/4 GlyT2^+^ vs. 8 Dbx1^+^: p=0.9; 8 Dbx1^+^/50 ms/8 GlyT2^+^ vs. 8 Dbx1^+^: p=0.3; 8 Dbx1^+^/100 ms/4 GlyT2^+^ vs. 8 Dbx1^+^: p=0.004; 8 Dbx1^+^/100 ms/8 GlyT2^+^ vs. 8 Dbx1^+^: p=0.004; n=9; Figure 3A-C, E). P_stim_ and amplitude and duration of evoked bursts were not significantly affected by 4 or 8 GlyT2^+^ neuron stimulation at 50 or 100 ms intervals (Friedman test, P_stim_: χ^2^(4)=0.6, p=1; amp: χ^2^(4)=1.51, p=0.8; T_I_: χ^2^(4)=5.8, p=0.2; n=9; Figure 3B, C, E). We increased the delay between 8 Dbx1^+^ neuron stimulation and either 4 or 8 GlyT2^+^ neuron stimulation to activate inhibitory neurons during the evoked burst (Figure 3D). GlyT2^+^ neuron stimulation during an evoked burst did not significantly alter burst amplitude or duration (Wilcoxon signed rank test, 4 GlyT2^+^ amp: p=0.4; 4 GlyT2^+^ T_I_: p=0.3; 8 GlyT2^+^ amp: p=0.1; 8 GlyT2^+^ T_I_: p=0.9; n=9; Figure 3D, E). Thus, stimulating 4 or 8 preBötC GlyT2^+^ neurons specifically delays burst initiation.

### Threshold preBötC GAD1^+^ stimulation prolongs burst duration

The ability to target specific numbers of neurons allowed us to directly compare whether the smaller subpopulation of preBötC GAD1^+^ neurons differed functionally from preBötC GlyT2^+^ neurons where GAD1 expression was largely absent. To assess the functional role of the GAD1^+^ preBötC subpopulation and specifically target these neurons, we used holographic stimulation in rhythmic medullary slices from Dbx1^tdTomato^;GAD1^EGFP^ (Dbx1^cre^ x Rosa26^LoxP-Stop-LoxP-tdTomato^ x GAD1^EGFP^) double reporter mice. Glutamate uncaging of a 10 μm soma-centered spot over a single preBötC GAD1^+^ neuron adjacent to Dbx1^+^ neurons resulted in a burst of action potentials (Figure 4A), similar to the responses of Dbx1^+^ and GlyT2^+^ neurons to photostimulation (Figure 2A; Kam et al., 2013b; Sun et al., 2019). We again confirmed the location of the field in preBötC by examining colocalization of Dbx1^+^ and GAD1^+^ and determining whether photostimulating 8 Dbx1^+^ neurons entrained inspiratory rhythm. We did not observe co-expression of Dbx1 and GAD1 in our target fields (Figure 4B). Photostimulation of 8 Dbx1^+^ neurons produced robust entrainment of the rhythm (P_stim_: 89.5% [IQR 80-97.2%]; n=7; Figure 4C), consistent with the field being in preBötC.

**Figure 4.**
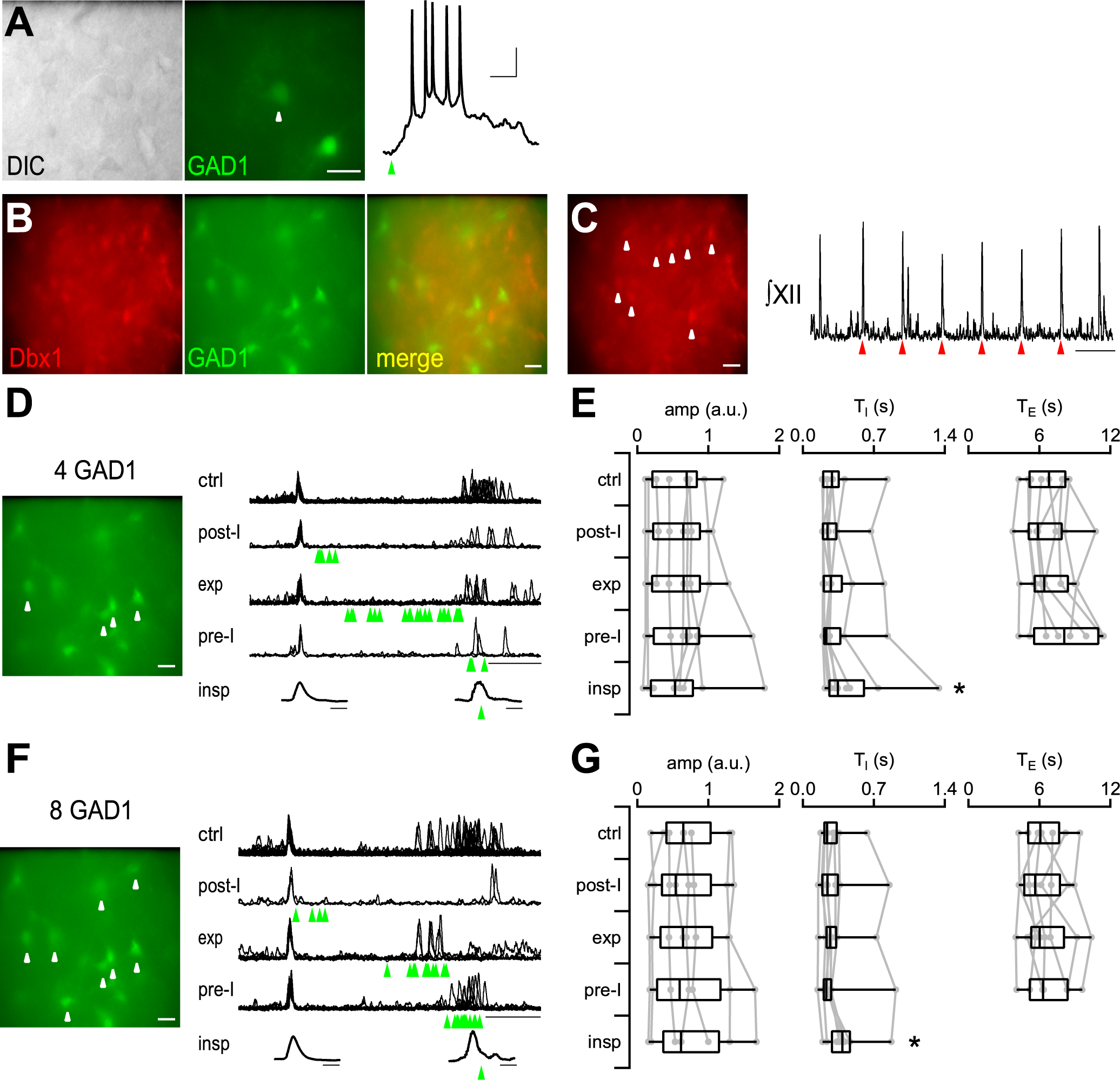
Stimulation of preBötC GAD1^+^ neurons during endogenous bursts increases T_I_. **(A)** DIC and fluorescence images of a Dbx1^tdTomato^;GAD1^EGFP^ mouse. Whole cell current clamp response of a GAD1^+^ neuron (white arrow) to photostimulation (green arrow). Scale bar, 10 mV, 100 ms. **(B)** Fluorescent images of preBötC from Dbx1^tdTomato^;GAD1^EGFP^ mouse. Scale bar, 100 µm. **(C)** *left*, Fluorescent tdTomato image of preBötC. White arrows mark targeted neurons. Scale bar, 100 µm. *right*, Integrated XII (∫XII) trace shows response to photostimulation of 8 Dbx1^+^ neurons. Scale bar, 5 s. **(D)** *left*, Fluorescent image showing 4 preBötC GAD1^+^ neurons targeted for photostimulation (white arrows). Scale bar, 100 µm. *right*, Overlay of cycles of ∫XII activity under control conditions with no stimulation, and when stimulation occurred during post-inspiratory (post-I), expiratory (exp), and preinspiratory (pre-I) phases. Green arrows mark time of stimulation. Scale bar, 2 s. Average control inspiratory burst shape (*bottom left*) and burst shape when stimulation occurred (*bottom right*) during the inspiratory burst (insp). Green arrow marks time of stimulation. Scale bar, 0.2 s. **(E)** Grouped amplitude (amp), burst duration (T_I_), and interburst interval (T_E_) for cycles with no stimulation (ctrl), and when stimulation of 4 GAD1^+^ preBötC neurons occurred during post-I, exp, and pre-I. T_I_ and amp are also shown when stimulation occurred during bursts (insp). *, p<0.05, Friedman test, post-hoc Wilcoxon signed rank test with Bonferroni-Holm correction, n=7 for post-I, exp, and pre-I, n=8 for insp. (**F)** *left*, Fluorescent image showing 8 preBötC GlyT2^+^ neurons targeted for photostimulation (white arrows). Scale bar, 100 µm. *right*, Overlay of cycles of ∫XII activity under control conditions with no stimulation, and when stimulation occurred during post-inspiratory (post-I), expiratory (exp), and preinspiratory (pre-I) phases. Green arrows mark time of stimulation. Scale bar, 2 s. Average control inspiratory burst shape (*bottom left*) and burst shape when stimulation occurred (*bottom right*) during the inspiratory burst (insp). Green arrow marks time of stimulation. Scale bar, 0.2 s. **(G)** Grouped amplitude (amp), burst duration (T_I_), and interburst interval (T_E_) for cycles with no stimulation (ctrl), and when stimulation of 8 GlyT2^+^ preBötC neurons occurred during post-I, exp, and pre-I. T_I_ and amp are also shown when stimulation occurred during bursts (insp). *, p<0.05, Friedman test, post-hoc Wilcoxon signed rank test with Bonferroni-Holm correction, n=7 for post-I, exp, and pre-I, n=8 for insp.

Following the protocol for preBötC GlyT2^+^ functional studies, we photoactivated 4 or 8 GAD1^+^ neurons during endogenous rhythm. In contrast to GlyT2^+^ stimulation, there were no statistically significant effects of the number of GAD1^+^ neurons stimulated or the phase on T_E_ (Friedman test, 4 GAD1^+^: χ^2^(3)=6.4, p=0.09; 8 GAD1^+^: χ^2^(3)=2.3, p=0.5; n=7; Figure 4D-G). The amplitude and duration of the burst following stimulation of 4 or 8 GAD1^+^ neurons at any phase was also not significantly different (Friedman test, 4 GAD1^+^ amp: χ^2^(3)=0.8, p=0.9; 4 GAD1^+^ T_I_: χ^2^(3)=1.5, p=0.7; 8 GAD1^+^ amp: χ^2^(3)=3.5, p=0.3; 8 GAD1^+^ T_I_: χ^2^(3)=3.5, p=0.3; n=7; Figure 4D-G). In contrast to GlyT2^+^ stimulation, photostimulation of 4 or 8 GAD1^+^ neurons during inspiratory bursts significantly increased burst duration by 21.2% (ctrl: 0.29 s [IQR 0.20-0.35 s]; 4 GAD1^+^: 0.35 s [IQR 0.26-0.60 s]) and 63.5% (ctrl: 0.24 s [IQR 0.21-0.33 s]; 8 GAD1^+^: 0.39 s [IQR 0.29-0.46 s]), respectively, while amplitude was not significantly affected (Wilcoxon signed rank test, 4 GAD1^+^ vs ctrl amp: p=0.2; 4 GAD1^+^ vs ctrl T_I_: p=0.04; 8 GAD1^+^ vs ctrl amp: p=0.7; 8 GAD1^+^ vs ctrl T_I_: p=0.02; n=8; Figure 4D-G)

We next determined whether GAD1^+^ stimulation could affect inspiratory rhythm and pattern generation using sequential stimulation. Photostimulation of 8 Dbx1^+^ neurons in slices from neonatal Dbx1^tdTomato^;GAD1^EGFP^ mice recapitulated the effects on P_stim_ and latency observed with Dbx1^tdTomato^;GlyT2^EGFP^ reporter mice (Figure 5A, E). When Dbx1^+^ stimulation was followed by stimulation of 4 or 8 GAD1^+^ neurons with a 50 or 100 ms delay, no significant differences in P_stim_, latency, burst amplitude, or burst duration were observed (Friedman test, P_stim_: χ^2^(4)=4.3, p=0.4; latency: χ^2^(4)=6.9, p=0.1; amp: χ^2^(4)=12.7, p=0.01; T_I_: χ^2^(4)=2, p=0.7; post-hoc Wilcoxon signed rank test with Bonferroni-Holm correction, amp: p>0.05 all comparisons; n=6; Figure 5A-C, E). To determine the effect of GAD1^+^ stimulation during the evoked burst, we increased the delay following 8 Dbx1^+^ stimulation. Stimulating 4 or 8 GAD1^+^ neurons during bursts evoked by stimulating 8 Dbx1^+^ neurons prolonged burst duration by 59% and 93%, respectively (ctrl: 0.24 s [IQR 0.21-0.32 s], n=6; 4 GAD1^+^: 0.39 s [IQR 0.30-0.50 s], n=5; 8 GAD1^+^: 0.47 s [IQR 0.26-0.59 s], n=6; Wilcoxon signed rank test, ctrl vs. 4 GlyT2^+^ T_I_: p=0.03; ctrl vs. 8 GlyT2^+^ T_I_: p=0.03; Figure 5D, E). While stimulating 4 GAD1^+^ neurons during evoked bursts did not affect burst amplitude, stimulating 8 GAD1^+^ neurons during evoked bursts caused a statistically significant 38.5% decrease in amplitude (ctrl: 6.9 a.u. [IQR 3.6-9.4 a.u.], n=6; 8 GAD1^+^: 4.2 a.u. [IQR 1.7-7.9 a.u.], n=6; Wilcoxon signed rank test, ctrl vs. 4 GAD1^+^ amp: p=0.4; ctrl vs. 8 GAD1^+^ amp: p=0.03; Figure 5D, E). We therefore demonstrate that stimulation of 4 or 8 preBötC GAD1^+^ neurons during endogenous and evoked bursts increases burst duration and stimulation of 8 preBötC GAD1^+^ neurons during evoked bursts decreases burst amplitude, revealing distinct functional roles for preBötC GlyT2^+^ and GAD1^+^ neurons.

**Figure 5.**
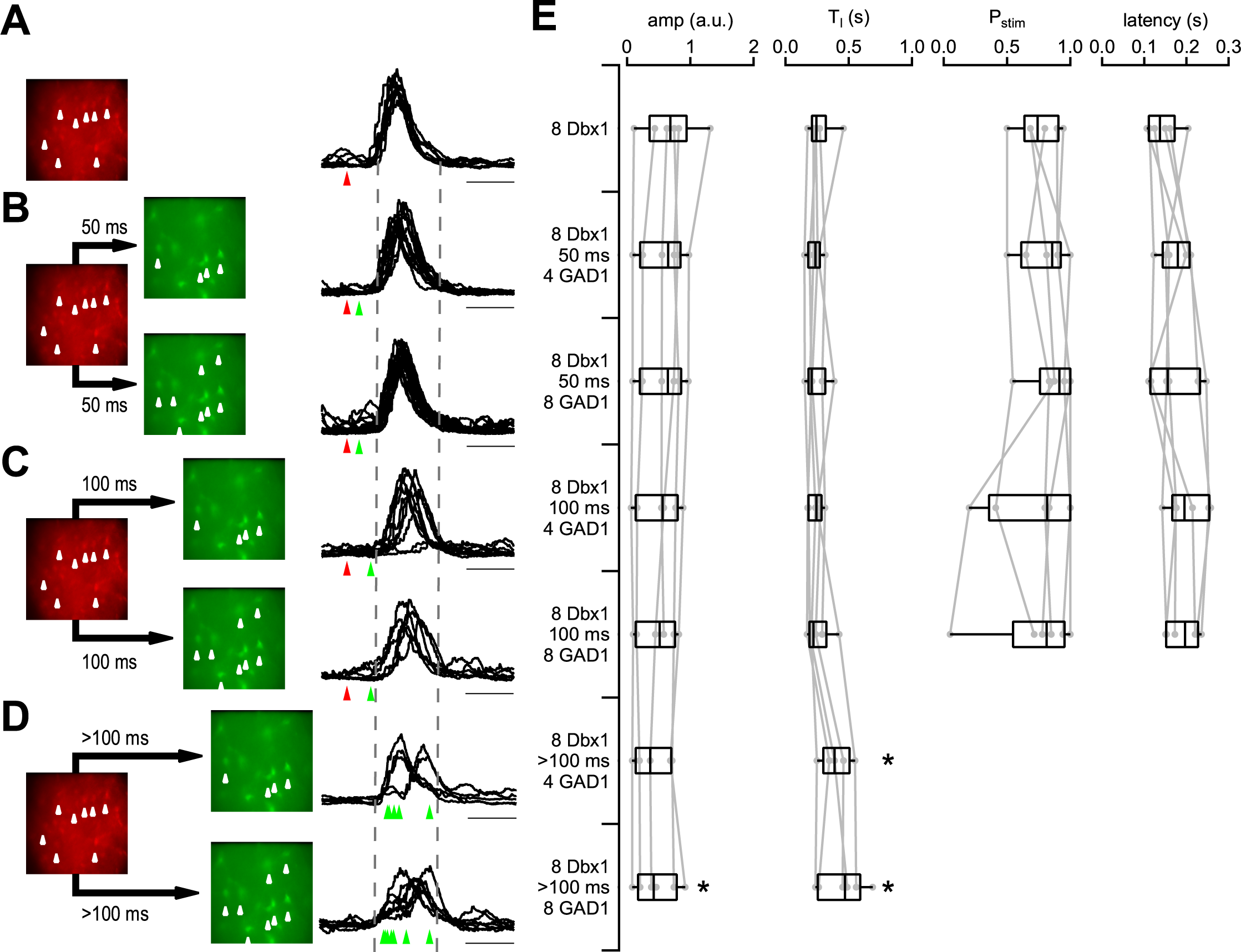
Stimulation of preBötC GAD1^+^ neurons during evoked bursts alters burst pattern. **(A-D)** *left*, Fluorescent images of preBötC from a Dbx1^tdTomato^;GAD1^GFP^ mouse and, *right*, overlaid traces from the ∫XII recording of evoked bursts triggered by **(A)** 8 Dbx1^+^ neurons, **(B)** 8 Dbx1^+^ neurons followed by 4 or 8 GAD1^+^ neurons after a 50 ms delay, **(C)** 8 Dbx1^+^ neurons followed by 4 or 8 GAD1^+^ neurons after a 100 ms delay, and **(D)** 8 Dbx1^+^ neurons followed by 4 or 8 GAD1^+^ neurons after a >100 ms delay (during the evoked burst). White arrows indicate targeted neurons, Red arrows indicate time of 8 Dbx1^+^ stimulation, Green arrows indicate time of 4 or 8 GAD1^+^ stimulation 50 ms, 100, ms, and > 100 ms after Dbx1^+^ stimulation. Dotted lines indicate average beginning and end of burst evoked by 8 Dbx1^+^ neurons. Scale bar, 0.2 s. **(E)** Grouped data showing amplitude (amp), burst duration (T_I_), probability of stimulation (P_stim_), and latency to burst initiation following stimulation of Dbx1^+^ and GAD1^+^ neurons. *, p<0.05, Wilcoxon signed rank test, n=5 8 Dbx1^+^/>100 ms/4 GAD1^+^, n=6 all other groups.

## Discussion

Inhibition appears to play several roles in modulation of breathing, but how these functions are governed by inhibitory neurons in the preBötC has not been clearly determined (Abdala et al., 2015; Ashhad et al., 2022; Dick et al., 2018; Feldman and Kam, 2015; Ramirez and Baertsch, 2018). We characterized the distribution of neurons expressing and co-expressing the major inhibitory markers GlyT2, GAD1, and GAD2 in preBötC. We determined that preBötC contains two major molecularly-defined inhibitory subpopulations in neonates, delineated by the presence or absence of GAD1. We then used holographic photostimulation to both establish that the neurons were in preBötC and to determine their functional role in endogenous rhythm and in evoked burst-generating processes. We found that GlyT2^+^ stimulation in the preinspiratory phase resulted in prolongation of the period and analogously that GlyT2^+^ photostimulation following threshold Dbx1^+^ stimulation resulted in increases in latency to burst initiation. In contrast to GlyT2^+^ stimulation, photostimulation of the same number of GAD1^+^ neurons did not delay endogenous or evoked inspiratory burst generation, but did prolong burst duration when stimulation occurred during endogenous or evoked inspiratory bursts. Activation of more GAD1^+^ neurons during inspiration decreased evoked burst amplitude. These results point to distinct functional roles for preBötC GAD1^+^ and GAD1^-^ neurons in the breathing central pattern generator.

The molecular heterogeneity of inhibitory preBötC neurons is consistent with previous studies examining numbers of molecularly-defined inhibitory neurons in preBötC and extend our understanding of the spatial distribution and molecular expression of these neurons. Using double reporter Dbx1^tdTomato^;GlyT2^EGFP^ mice to visualize Dbx1^+^ and GlyT2^+^ populations in the same tissue, we found approximately equal numbers of Dbx1^+^ and GlyT2^+^ neurons. In previous studies using GlyT2^EGFP^ mice, GlyT2^+^ neurons were found to constitute approximately half of inspiratory-modulated neurons in preBötC, with some having intrinsic bursting properties (Morgado-Valle et al., 2010; Winter et al., 2009). We found more overlap between Dbx1^+^ and GlyT2^+^ neurons in our histological analysis than previously reported (Bouvier et al., 2010; Gray et al., 2010; Morgado-Valle et al., 2010; Winter et al., 2009); although, we did not see significant overlap in the acute slices. GlyT2, GAD1, and GAD2 transcripts were detected in Dbx1^+^ preBötC neurons (Hayes et al., 2017), so preBötC Dbx1^+^ neurons may transiently express GlyT2; our analysis may have included neurons outside the ventral respiratory column and preBötC as defined in previous studies; and/or there may be some non-specific cre-expression in the Dbx1^cre^ mouse. We found similar numbers of GlyT2^+^ and GAD2^+^ neurons in preBötC, perhaps due to their substantial co-expression, consistent with previous studies (Hirrlinger et al., 2019; Oke et al., 2023). While GlyT2^+^ and GAD2^+^ neurons constituted a large number of preBötC neurons, there were significantly fewer GAD1^+^ neurons in preBötC.

Quantitative differences in our results and those reported in other studies may be due to different boundaries for preBötC and, in some cases, a focus on inspiratory-modulated neurons. preBötC, like many functionally-defined structures in the reticular formation, does not have clear anatomical borders, despite the position of these structures being highly stereotyped (Feldman and Kam, 2015). As neither Dbx1, nor any of the inhibitory markers, delineate preBötC boundaries, we opted to use a hemicylindrical volume that hews roughly to a functionally defined rostrocaudal extent 360-480 µm caudal to caudal pole of facial motor nucleus and a transverse area corresponding roughly to the lateral part of the paragigantocellular reticular nucleus or the medial part of the lateral reticular nucleus (Ruangkittisakul et al., 2014).

Molecular co-expression of GlyT2, GAD2, and GAD1 further highlights the heterogeneity of preBötC inhibitory neurons. In our study, GlyT2 and GAD2 were frequently co-expressed in preBötC neurons. GAD1, where expressed, also overlapped frequently with both GlyT2 and GAD2. The high degree of overlap between GlyT2 and GAD2 points to a significant proportion of triple-expressing GAD1^+^/GAD2^+^/GlyT2^+^ preBötC inhibitory neurons. Using a split-Cre system to track expression of GlyT2 and GAD2, a significant number of preBötC neurons were found to express both GlyT2 and GAD2 in neonatal mice (Hirrlinger et al., 2019; Oke et al., 2023). GlyT2 and GAD1 co-expression was demonstrated with single cell multiplex RT-PCR of samples from GAD1^+^ neurons and GlyT2 immunostaining in GAD1^EGFP^ mice (Koizumi et al., 2013). While GAD2 and GAD1 are frequently co-expressed in cortical GABAergic neurons (Erlander et al., 1991; Soghomonian and Martin, 1998), we find that GAD1/GAD2 co-expression is more limited in neonatal preBötC. In other neurons, GAD2 has been found to localize to synaptic terminals while GAD1 appears concentrated in the somatic compartment (Erlander et al., 1991; Soghomonian and Martin, 1998). GAD2 may therefore serve to replenish the neurotransmitter pool of GABA while GAD1 may play a metabolic role (Erlander et al., 1991; Soghomonian and Martin, 1998). Whether the GAD isoform or GABA-glycine co-release plays a role in the functional differences remains to be determined. Co-expression across these inhibitory markers was not complete, so singly expressing neurons or neurons expressing other combinations, such as GAD1^+^/GAD2^+^ or GAD1^+^/GlyT2^+^ double expressing neurons, could constitute other functionally distinct subpopulations. Co-expression of GlyT2 and GAD2 is developmentally regulated, with many preBötC inhibitory neurons expressing only GlyT2 or GAD2 in adults (Hirrlinger et al., 2019; Oke et al., 2023). Whether GlyT2 or GAD2 is specifically developmentally downregulated in GAD1^+^ neurons or whether GAD1 is also downregulated during maturation remains to be determined. However, in neonatal mice, we conclude that there are two major inhibitory subpopulations: neurons co-expressing GAD2 and GlyT2, and those expressing GAD2, GAD1, and GlyT2.

We aimed to determine whether preBötC GAD1^+^ and GAD1^-^ inhibitory populations play functionally distinct roles in inspiratory rhythm and pattern generation *in vitro*. Pharmacological blockade of inhibitory transmission, local microinjection, and opsin expression induced in whole animal transgenic mice or by viral infection often affect nearby BötC neurons, and, in the case of whole animal opsin expression, inhibitory inputs from afferents throughout the brainstem (Ausborn et al., 2018; Baertsch et al., 2018; Fortuna et al., 2019; Huff et al., 2022; Hulsmann et al., 2021; Sherman et al., 2015; Vafadari et al., 2023). To overcome this technical obstacle, we used holographic photostimulation to restrict activation to a small number of neurons and to establish that the targeted inhibitory neurons were in preBötC by showing that activating adjacent Dbx1^+^ neurons entrained inspiratory rhythm.

We found that photostimulating 4 or 8 GlyT2^+^ preBötC neurons during the preinspiratory period produced a phase delay. These data are consistent with brief optogenetic activation of GlyT2^+^ neurons causing phase-dependent delays in inspiratory activity *in vivo* with phase delays being strongest when stimulation occurred in the preinspiratory period (Sherman et al., 2015). Prolonged stimulation extends the expiratory period and can produce apnea (Fortuna et al., 2019; Sherman et al., 2015). Repeated optogenetic stimulation may also terminate inspiration or increase breathing frequency by eliciting rebound excitation (Baertsch et al., 2018; Fortuna et al., 2019; Hulsmann et al., 2021). Whether these other effects are due to activation of a greater number of preBötC GlyT2^+^ neurons; stimulation of GlyT2^+^ neurons in preBötC, BötC, and inhibitory afferents farther away; and/or indiscriminate activation of multiple inhibitory subpopulations due to the use of the vesicular GABA transporter (VGAT) promoter remains to be tested.

We also found that activation of 4 or 8 preBötC GAD1^+^ neurons unexpectedly increased burst duration when stimulation occurred during the inspiratory burst and that stimulation of 8 GAD1^+^ neurons also decreased evoked burst amplitude. Previous studies pointed to a role for inhibitory neurons in decreasing burst amplitude and duration. Pharmacological blockade of GABA_A_ receptors increased amplitude and modestly increased duration (Baertsch et al., 2018; Janczewski et al., 2013; Shao and Feldman, 1997). Optogenetic GlyT2^+^ and VGAT^+^ neuron stimulation *in vivo* decreased inspiratory duration or terminated inspiration (Baertsch et al., 2018; Hulsmann et al., 2021; Sherman et al., 2015). However, these studies did not specifically activate GAD1^+^ preBötC neurons. Our data are thus the first to show how targeted activation of GAD1^+^ preBötC neurons affects inspiratory rhythm and pattern generation.

While these threshold effects were statistically significant, there was large variability in responses to stimulation that may have diminished effects sizes. Some of the variability may be explained by further molecular and functional heterogeneity within preBötC GAD1^+^ and GAD1^-^ inhibitory subpopulations (Koizumi et al., 2013; Zheng et al., 2019). Intrinsic properties of inhibitory neurons expressing GAD1, GlyT2, or both do not appear to differ significantly across these subpopulations, but do show some differences with excitatory inspiratory neurons (Burgraff et al., 2022; Koizumi et al., 2013; Picardo et al., 2013; Picardo et al., 2019; Reising et al., 2022; Zheng et al., 2019). Other sources of variability that may dilute the effects of photostimulation include differences in slice preparations and in single neuron responses to photoactivation. Some GlyT2^+^ neurons could also be GAD1^+^ neurons, and there is a high probability that GAD1^+^ neurons expressed GlyT2^+^. Nonetheless, the probability of stimulated GlyT2^+^ neurons also being GAD1^+^ is quite small, and the differential effects of stimulating GlyT2^+^ vs. GAD1^+^ support distinct functionality between these two subpopulations in spite of the small number of neurons activated and numerous potential sources of variability.

The effects of stimulating a small number of neurons have revealed fundamental properties of neural circuits in mammals (Doron and Brecht, 2015) and provide unique constraints for computational models of preBötC (Feldman and Kam, 2015). Single primary motor cortical neuron stimulation can initiate whisking, demonstrating how modulation of one or a few neurons can alter a rhythmic motor behavior (Brecht et al., 2004). Additionally, these threshold effects, particularly threshold numbers of neurons, are critical network parameters that affect neural circuit dynamics (Feldman and Kam, 2015). The 4-9 inspiratory preBötC neuron threshold for triggering an evoked burst *in vitro* has served as a useful constraint for computational models that point to experimentally testable network parameters and mechanisms (Ashhad et al., 2023; Guerrier et al., 2015; Phillips and Rubin, 2022; Song et al., 2015). Indeed, the cell number-dependent effects on evoked burst amplitude with GAD1^+^ neuron stimulation may suggest that regulation of distinct aspects of inspiratory rhythm and pattern generation may be achieved by tuning the number of active preBötC inhibitory neurons. These data may therefore constrain computational models that include inhibitory preBötC neurons and point to novel mechanisms of inhibitory regulation of breathing.

While conceptually useful, are the threshold effects of stimulating GlyT2^+^ and GAD1^+^ preBötC neurons physiologically relevant? Entire classes of inhibitory neurons are often assumed to fire synchronously (Kvitsiani et al., 2013) even though activation of a few, or even single, neurons can significantly impact neural circuit dynamics and subserve functions other than blanket inhibition (Isaacson and Scanziani, 2011; Karnani et al., 2016a; Karnani et al., 2016b). Indeed, a major role for inhibition is in regulating excitation-inhibition balance where even minor alterations in the number or strength of inhibitory activity can alter neural circuit dynamics (Ashhad and Feldman, 2020; Isaacson and Scanziani, 2011; Ramirez and Baertsch, 2018). The overlapping connectivity and phasic activity of inhibitory and excitatory preBötC neurons supports such a role. Excitatory SST^+^ and GlyT2^+^ preBötC neurons overlap extensively in their efferent targets as well as in their inputs (Yang and Feldman, 2018; Yang et al., 2020). Morphological analysis of GAD1^+^ and GlyT2^+^ inspiratory preBötC neurons also showed overlap with glutamatergic preBötC neurons in their projections dorsomedially to XII premotoneuron and motoneuron areas as well as in their dendritic fields (Koizumi et al., 2013). GlyT2^+^ and VGAT^+^ neurons have been found to be tonically active, expiratory-modulated, and inspiratory-modulated, some with intrinsic bursting properties (Baertsch et al., 2018; Burgraff et al., 2022; Ezure et al., 2003; Morgado-Valle et al., 2010; Oke et al., 2018; Winter et al., 2009). Similarly, the majority of GAD1^+^ neurons also displayed inspiratory-modulated activity (Koizumi et al., 2013), suggesting that GlyT2^+^ and GAD1^+^ preBötC neurons co-terminate and are co-active with excitatory Dbx1^+^ preBötC neurons during preinspiration and inspiration.

These data support a model whereby GlyT2^+^ and GAD1^+^ preBötC neurons contribute to excitation-inhibition balance governing distinct respiratory-related processes: GlyT2^+^ to burst initiation and inspiratory rhythmogenesis and GAD1^+^ to burst duration and amplitude and inspiratory patterning. While inhibition is not necessary for inspiratory rhythmogenesis (Janczewski et al., 2013; Shao and Feldman, 1997; Sherman et al., 2015), preBötC GlyT2^+^ activity may play a role in reducing cycle to cycle variability (Hirrlinger et al., 2019), and changes in GlyT2^+^ preBötC neuronal activity during preinspiration may allow precise, rapid regulation of burst timing by advancing or delaying bursts when GlyT2^+^ neuron activity is decreased or increased (Baertsch et al., 2018; Sherman et al., 2015). GAD1^+^ preBötC neuron activity during inspiration, on the other hand, modulates excitation-inhibition balance onto preBötC pattern generating neurons, inspiratory premotoneurons, and/or XII motoneurons to increase or decrease burst duration and amplitude. We suggest that differences in connectivity, phasic firing properties, and/or levels of excitation and inhibition during the interburst interval compared to levels of excitation and inhibition during inspiratory bursts may explain how GlyT2^+^ has an inhibitory effect on burst initiation while GAD1^+^ appears to have an excitatory effect on burst duration. Activation of more preBötC GlyT2^+^ or GAD1^+^ neurons, as observed in optogenetic studies, may shift the excitation-inhibition balance towards inhibition and result in stronger inhibitory effects on both rhythm and pattern even when stimulation occurs outside of the preinspiratory or inspiratory period.

In summary, we discover functionally distinct inhibitory preBötC subpopulations defined by the presence or absence of GAD1. We find that threshold stimulation can precisely and rapidly regulate the initiation of inspiration through activation of GlyT2^+^ neurons and the duration and magnitude of inspiration through activation of GAD1^+^ neurons. We suggest that these modest changes in GlyT2^+^ and GAD1^+^ preBötC activity alter excitation-inhibition balance in rhythm and pattern generating microcircuits, allowing for precise, dynamic control of breathing. We conclude that the activity of distinct subpopulations of inhibitory preBötC neurons constitute functional microcircuits that can rapidly shape breathing, expand dynamic range, and confer lability that allows breathing to respond to changes in metabolic demand and environmental conditions and to coordinate with other respiratory-related behaviors, such as swallowing and vocalization.

## Acknowledgements

GlyT2^cre^ and GlyT2^EGFP^ mice were generously provided by Dr. H.U. Zeilhofer. This work was supported by National Institutes of Health grant NS097492.

## Competing Interests statement

The authors declare no competing interests.

## Materials and methods

Experimental procedures were carried out in accordance with the United States Public Health Service and Institute for Laboratory Animal Research Guide for the Care and Use of Laboratory Animals. All animals were handled according to approved institutional protocols at Rosalind Franklin University of Medicine and Science (#B14-16, #B18-10, #B21-03). All protocols were approved by the Rosalind Franklin University of Medicine and Science Institutional Animal Care and Use Committee (Animal Welfare Assurance #A3279-01). Animals were subject to regular veterinary inspection and only to the experimental procedures detailed here and maintained on a conventional 12hr light/12hr dark cycle with free access to food and water. Every effort was made to minimize pain and discomfort.

### Transgenic mice

Rosa26^LoxP-Stop-LoxP-tdTomato^ (JAX Stock No. 7914; Madisen et al., 2010) mice were crossed with either GlyT2^EGFP^ (Zeilhofer et al., 2005) or GAD1^EGFP^ (JAX Stock No. 7677; Chattopadhyaya et al., 2004) BAC transgenic mice to generate Rosa26^LoxP-Stop-LoxP-^tdTomato;GlyT2^EGFP^ and Rosa26^LoxP-Stop-LoxP-tdTomato^;GAD1^EGFP^ mouse lines. These mice were then crossed with either Dbx1^cre^ (Bielle et al., 2005), GlyT2^cre^ (Foster et al., 2015), or GAD2^cre^ (JAX Stock No. 19022; Taniguchi et al., 2011) mice to generate Dbx1^tdTomato^;GlyT2^EGFP^, Dbx1^tdTomato^;GAD1^EGFP^, GAD2^tdTomato^;GlyT2^EGFP^, GAD2^tdTomato^;GAD1^EGFP^, and GlyT2^tdTomato^;GAD1^EGFP^ double reporter mice. Animal genotypes were verified via real time polymerase chain reaction using primers specific for cre recombinase or by direct visualization of fluorescent reporter.

### Histology and imaging

Male and female neonatal double reporter mice (P0-P4) were anesthetized by inhalation of isoflurane and transcardially perfused with 4% paraformaldehyde. A block of tissue from the pons to the rostral cervical spinal cord was isolated, fixed overnight in 4% paraformaldehyde, and cryoprotected in 30% sucrose. The fixed brainstem-spinal cord block was sliced coronally on a Leica CM3050S cryostat in 60 μm intervals from the obex to the superior vestibular nucleus.

Sections were cover-slipped in PVA-DABCO (Valnes and Brandtzaeg, 1985) and imaged on a confocal Leica SP8 microscope. Facial motor nucleus and nucleus ambiguus (NA) were identified by the absence of GlyT2^+^, GAD1^+^, and GAD2^+^ neurons in reporter mice and used as landmarks to identify preBötC. For each section, the preBötC was centered within a 450 μm × 450 μm field and a z-stack (1.5 μm optical section) obtained using a 20x objective.

Z-stacks were imported into CellProfiler 4.2.1 (Carpenter et al., 2006) one channel at a time and run through a custom-designed image processing pipeline to automatically extract the total cell count and 3-dimensional coordinates of cell centroids. Briefly, image intensity was first rescaled to utilize the full dynamic range. Images were then shrunk to half their size and spatially averaged to reduce over-segmentation and increase the uniformity of signal within labeled cells. Resized images were subject to spatial averaging and morphological operations to enhance soma shapes. A sequential threshold was then applied, followed by further morphological operations to remove artifacts and fill holes. A watershed algorithm was then used to segment cells across three dimensions. Image processing parameters were determined empirically for each channel. Fluorescence intensity, shape, and position of detected cells were then measured and written to an output file. Image processing pipelines are available from the corresponding author’s website (https://sites.google.com/site/kwkamlab) or the CellProfiler pipeline repository (https://cellprofiler.org/published-pipelines).

CellProfiler output and confocal image stacks were loaded into IgorPro for manual review and editing. Custom written code was utilized to overlay the coordinate information extracted from CellProfiler onto the raw images from each channel. A 3-dimensional distance threshold was then applied to determine whether neurons expressed both markers, i.e., were co-expressing. Cells within a 150 µm semicircle centered at the ventral pole of NA were counted, and counts from left and right sides were summed for each rostrocaudal section.

### Slice electrophysiology

Neonatal double reporter mice (P0-P4) of either sex were anesthetized with isoflurane and euthanized by thoracic transection. Brainstems were removed in cold artificial CSF (ACSF) containing the following (in mM): 124 NaCl, 3 KCl, 1.5 CaCl_2_, 1 MgSO_4_, 25 NaHCO_3_, 0.5 NaH_2_PO_4_, and 30 glucose, which we aerated with 95% O2 and 5% CO2. Brainstems were then pinned to an agar block with the rostral side up. We cut a single 550-600 μm thick transverse medullary slice with the preBötC at the rostral surface. To obtain slices with the preBötC at the surface, the rostral cut was made above the first set of XII nerve rootlets at the level of the dorsomedial cell column and principal lateral loop of the inferior olive, and the caudal cut captured the obex (Ruangkittisakul et al., 2014). Slices were placed rostral side up in a chamber and perfused at 2– 3 ml/min with 28–30°;C recording ACSF. Baseline recording ACSF consisted of the cutting ACSF solution with K^+^ raised to 9 mM and Ca^2+^ maintained at 1.5 mM for a robust burst rhythm (Kam et al., 2013a). In all experiments, slices were allowed to equilibrate for 30 min to ensure that the frequency and magnitude of XII population activity reached steady-state. Respiratory activity reflecting suprathreshold action potential firing from populations of neurons was recorded from XII motor nucleus or nerve roots or as population activity directly from the preBötC using suction electrodes (tip size ∼50 µm). For whole cell patch clamp recordings, preBötC GlyT2^+^ or GAD1^+^ neurons were selected visually using fluorescence microscopy and patched under IR-DIC video microscopy. Patch electrodes with a 2–3 M tip resistance were fabricated using a three-stage custom program on a Flaming-Brown micropipette puller (P-97, Sutter Instruments) and filled with solution containing the following (in mM): 135 KGluconate, 1.1 EGTA, 5 NaCl, 0.1 CaCl_2_, 10 HEPES, 2 MgATP, and 0.3 Na_3_GTP. In whole-cell current-clamp recordings, membrane potentials were brought to −55 mV using a negative bias current. For whole-cell recordings, experiments in which variations in series and input resistance exceeded the initial value by 25% were discarded. Population and single cell recordings were amplified using a Multiclamp 700B (Molecular Devices), filtered at 2–4 kHz, integrated, and digitized at 10 kHz. Integration was performed on custom built Paynter filter with a 20– 100 ms time constant. Digitized data were analyzed off-line using custom procedures written for IgorPro (Wavemetrics)

### Holography

Holographic photostimulation was performed on a Phasor SLM system (Intelligent Imaging Innovations, Inc) mounted around an epifluorescence upright microscope (Zeiss Axioscope). We used a 405 nm diode laser (CUBE 405–100; Coherent) to uncage MNI-glutamate (0.5 mM) and depolarize targeted neurons. An iterative Fourier transform algorithm, implemented in Slidebook 6 (Intelligent Imaging Innovations, Inc), computed the phase pattern on the SLM corresponding to the desired distribution of light intensity at the focal plane of the objective. A blocker was inserted in the intermediate Fourier plane to block the unmodulated light component (zero order spot) and replicate patterns (Golan et al., 2009). Neurons were targeted by centering 10 μm circular spots over the somata of the selected neurons. The laser intensity per spot was set at 1–3 mW, which was previously determined to elicit bursts of action potentials in the targeted neurons without the spread of glutamate to neighboring neurons (Kam et al., 2013b). Each laser stimulation consisted of 5 × 0.8 ms pulses, delivered at 200 Hz.

### Data analysis and statistics

Semiautomated event detection of respiratory-related activity recorded in XII output or preBötC population recordings was performed using custom procedures written in IgorPro. Multiple criteria, including slope and amplitude thresholds, were used to select events automatically, which were then confirmed visually.

Data are expressed as mean ± standard deviation or median [interquartile range (IQR)]. Box plots show median, IQR, and 10% and 90% thresholds. In figures displaying average data, data points from individual experiments are displayed in gray. To determine sample sizes, *a priori* power analysis was performed using G*power 3.1.9.2 (Faul et al., 2009). In this program, the appropriate statistical test along with a desired power of 95% at a 5% significance level was selected for each type of experiment with effects sizes based on published or preliminary data.

These parameters yielded minimum total sample sizes of 6 for all tests. Sample sizes represent the number of biological replicates for each group. Statistical analysis was performed using IgorPro. Data were tested for normality using the Shapiro Wilk test. One-way ANOVA was used for parametric multiple comparisons followed by a post-hoc Tukey test to determine statistical significance. For data that were not normally distributed, Wilcoxon signed rank tests were used for paired comparisons of two groups. Multiple comparisons were evaluated using Friedman test for repeated measures designs. Post-hoc Wilcoxon signed rank tests with Bonferroni-Holm corrections were then used to determine significance for within group comparisons of repeated measures designs. Statistical significance was set at p < 0.05.

## Notes

### Competing Interest Statement

The authors have declared no competing interest.

### Summary of Updates

Manuscript text and figures have been updated

